# New genetic insights about hybridization and population structure of hawksbill and loggerhead turtles from Brazil

**DOI:** 10.1101/2020.04.17.046623

**Authors:** Larissa S. Arantes, Sibelle T. Vilaça, Camila J. Mazzoni, Fabrício R. Santos

## Abstract

An extremely high incidence of hybridization among sea turtles is found along the Brazilian coast. To understand this atypical phenomenon and its impact on sea turtle conservation, research focused in the evolutionary history of sea turtles is fundamental. We assessed high quality multilocus haplotypes of 143 samples of the five species of sea turtles that occur along the Brazilian coast to investigate the hybridization process and the population structure of hawksbill (*Eretmochelys imbricata*) and loggerhead turtles (*Caretta caretta*). The multilocus data were initially used to characterize interspecific hybrids. Introgression (F2 hybrids) was only confirmed in hatchlings of F1 hybrid females (hawksbill × loggerhead), indicating that introgression was either previously overestimated and F2 hybrids may not survive to adulthood, or the first-generation hybrid females nesting in Brazil were born as recent as few decades ago. Phylogenetic analyses using nuclear markers recovered the mtDNA-based Indo-Pacific and Atlantic lineages for hawksbill turtles, demonstrating a deep genetic divergence dating from the early Pliocene. In addition, loggerhead turtles that share a common feeding area and belong to distinct Indo-Pacific and Atlantic mtDNA clades present no clear genetic differentiation at the nuclear level. Finally, our results indicate that hawksbill and loggerhead rookeries along the Brazilian coast are likely connected by male-mediated gene flow.

## Introduction

Sea turtles have complex life cycles, with life stages associated with different environments affected directly by human activities. This close interaction exposes them to several threats, which led to a global population decline of most species during the XX century. The main threats are related to fisheries bycatch, coastal urbanization, pollution (sewage, garbage, toxic substances), pathogens and exploitation of eggs, meat or other turtle products (Wallace et al. 2011). Thus, the monitoring of sea turtle populations followed by actions to mitigate the anthropogenic impact are essential to their conservation worldwide.

Understanding the population dynamics of sea turtles is challenging due to their highly migratory behavior, long lives and different levels of population structure associated with each life stage (Bowen and Karl 2007). Methodological advances in the last decades, such as satellite telemetry and molecular analyses, made important contributions to the comprehension of complex sea turtle behaviors, deepening the knowledge about the factors affecting the composition of foraging aggregations (Carreras et al. 2011; Proietti et al. 2014b), the migration route of pelagic juveniles also known as “lost years” (Putman and Mansfield 2015; Briscoe et al. 2016), the frequency of occurrence of multiple paternity (Moore and Ball 2002; González-Garza et al. 2015), the level of gene flow among populations (Bowen et al. 2005; Monzón-Argüello et al. 2011; Clusa et al. 2018) and opportunistic mating systems (Stewart and Dutton 2011). Novel technologies have also allowed to investigate the arguable reproductive isolation between sea turtle species, as interspecific hybridization was detected among five out of seven extant species of sea turtles (Vilaça et al. 2012). Some hybridization cases involve crosses of species of the Cheloniidae family that diverged at 63 million years ago (Naro-Maciel et al. 2008), probably the most deeply divergent species group capable of producing viable hybrids in nature (Karl et al. 1995).

Hybrid zones of sea turtles may occur where there is an overlap of nesting areas and reproductive seasons of two or more species, which occurs in two coastal areas of Brazil (Soares et al. 2017). The hybridization process of sea turtles in the northeastern Brazilian coast is atypical, since the frequency of hybrids is much higher than in any other analyzed population worldwide. While hybridization cases have been sporadically reported around the world (Karl et al. 1995), the frequency of hybrid females along the Brazilian coast reach frequencies as high as 42% in some nesting sites (Lara-Ruiz et al. 2006; Reis et al. 2010b).

In the northern coast of the Bahia state in Brazil, where the largest in-country rookeries of hawksbill (*Eretmochelys imbricata*) and loggerhead turtles (*Caretta caretta*) are found, 42% of female turtles morphologically identified as *E. imbricata* exhibited mitochondrial sequences of *C. caretta* (Lara-Ruiz et al. 2006). A more recent study confirmed that the incidence of hybrids is as high as 31.58% of the assumed *E. imbricata* population (Soares et al. 2018). The majority of the surveyed hybrids appears to have 50% of alleles of each parental species, thus being considered as first-generation (F1), but backcrossing with both parental species was also detected, revealing the occurrence of introgression (Vilaça et al. 2012). Because some backcrossed nesting females were also found, hybridization was estimated to have started at least two generations ago (>40 years), during a period when populations were heavily depleted due to the anthropogenic impact (Vilaça et al. 2012).

In a nearby nesting site in the Sergipe state of Brazil, where olive ridley (*Lepidochelys olivacea*) and loggerhead turtles present a spatial and temporal overlapping distribution, 27% of individuals morphologically identified as loggerheads were shown to be hybrids between both species (Reis et al. 2010b), all of them classified as F1 hybrids (Vilaça et al. 2012). This large frequency of interspecific hybrids in Brazilian rookeries is an important conservation concern because it may result in outbreeding depression, which is a decrease of the fitness and/or reproductive viability of local populations (Allendorf et al. 2001; Maheshwari and Barbash 2011). Outbreeding depression can be observed in F1 hybrids, and also in F2 or later generations due to the disruption of coadapted gene complexes as a result of meiotic recombination during gametogenesis in F1 hybrids (Goldberg et al. 2005). However, other studies suggest that interspecific hybridization may eventually represent an important source of variation as it may confer an advantageous effect on fitness, also called adaptive introgression (Hedrick 2013). In any case, it is extremely necessary to carefully investigate the consequences of hybridization for the populations where it occurs in high frequency, like the ones in Brazil.

Two previous studies (Soares et al. 2017, 2018) evaluated the potential outbreeding depression effects of hybridization via the comparison of several reproductive parameters between F1 hybrids and parental species in a nesting site located in Bahia. Even though emergence success was shown to be lower for hybrid nests, other parameters such as the hatchling production per clutch and clutch frequency were similar to parental species, suggesting that hybrids may persist in this region (Soares et al. 2017). The initial viability of hybrid hatchlings was also similar to non-hybrid hatchlings, revealing no significant evidence for hybrid breakdown at this early stage (Soares et al. 2018). However, once hybrid hatchlings achieve the sea, little is known about their survival until adulthood and reproductive fitness.

Other genetic studies including Brazilian sea turtles have investigated their demographic history (Bjorndal et al. 2006; Vargas et al. 2008; Molfetti et al. 2013), population structure (Reis et al. 2010a; Vilaça et al. 2013; Shamblin et al. 2014; Arantes et al. 2020), mixed stocks at foraging aggregations (Proietti et al. 2009; Reis et al. 2010a; Vilaça et al. 2013; Proietti et al. 2014b) and interspecific hybridization (Lara-Ruiz et al. 2006; Reis et al. 2010b; Vilaça et al. 2012; Proietti et al. 2014a; Soares et al. 2017, 2018). For example, phylogeographic analyses using mtDNA showed significant genetic divergence among three Brazilian rookeries of *C. caretta*, suggesting the recognition of three different management units (Shamblin et al. 2014), while two separate demographic units were recognized for *E. imbricata* in Brazilian nesting areas (Vilaça et al. 2013). The mixed stocks found at foraging aggregations along the Brazilian coast demonstrated connectivity among distant ocean basins (Reis et al. 2010a), which is highly influenced by oceanic currents (Vilaça et al. 2013; Proietti et al. 2014b).

All the above-cited studies have used data from mitochondrial DNA (mtDNA), microsatellites and/or other few nuclear (nDNA) markers, but recent advances in high-throughput sequencing technology (NGS) have opened up new opportunities to assess genome-wide data in a cost-effective way. Indeed, recent population genomic methods have allowed the survey of selected subsets of genetic markers in many individuals simultaneously (Harrisson et al. 2014). Furthermore, genome-wide multilocus data is being increasingly used in many ecological and evolutionary studies, providing inferences about the life history, population dynamics and demographic patterns of species, with important conservation implications (Davey et al. 2011; Harrison et al. 2014).

In this work, we designed a highly informative multilocus panel based on a genomic survey for the identification and characterization of interspecific hybrids between *E. imbricata* and *C. caretta*. Informative loci/haplotypes were selected from double-digest RADseq (ddRAD; Peterson et al. 2012) data produced for both species. For each selected locus, high-quality phased sequences were produced via Sanger sequencing and used for both characterization of hybrids and population studies. We compared our results with previous studies to test the efficiency of ddRAD-derived resequenced multilocus markers in increasing knowledge about the hybridization phenomenon and population structure of sea turtles along the Brazilian coast.

## Materials and Methods

### Sampling

We analyzed 143 DNA samples from five species of sea turtles and hybrid individuals from four different hybrid classes (Vilaça et al. 2012) that occur along the Brazilian coast (Table 1). The DNA samples were derived from individuals collected between 1999 and 2011 by the Projeto TAMAR team, a consolidated and successful Brazilian Sea Turtle Conservation Program. Some samples have already been surveyed in previous studies of our research group using mtDNA and few autosomal markers (Lara-Ruiz et al. 2006; Reis et al. 2010a; Vilaça et al. 2012; Vilaça et al. 2013). The species, localities and number of individuals analyzed are shown in Table 1. Detailed information of hybrid individuals (locality, morphology, collected individual, mtDNA haplotype based on control region and previous classification by Vilaça et al. 2012) is available in Supplementary Table S1. All *C. caretta* × *E. imbricata* (*Cc* × *Ei*), *E. imbricata* × *L. olivacea* (*Ei* × *Lo*) and *E. imbricata* × *C. caretta* × *Chelonia mydas* (*Ei* × *Cc* × *Cm*) hybrids were reported in Praia do Forte (Bahia) nesting site, and *C. caretta* × *L. olivacea* (*Cc* × *Lo*) hybrids were reported from Pirambu (Sergipe) nesting site. Other few hybrid individuals were reported from foraging aggregations or bycatch in fisheries.

**Table 1.** Sampling localities of sea turtles and hybrids and number of individuals per locality (N). *C. caretta* × *E. imbricata* (*Cc* × *Ei*), *E. imbricata* × *L. olivacea* (*Ei* × *Lo*), *C. caretta* × *L. olivacea* hybrids (*Cc* × *Lo*), *E. imbricata* × *C. caretta* × *C. mydas* (*Ei* × *Cc* × *Cm*).

| Sea turtle species | Sample Locality |  | N |
| --- | --- | --- | --- |
| **Loggerhead turtle**<br>(*Caretta caretta*) | Foraging Area | Elevação do Rio Grande | 15 |
|  | Nesting Area | Bahia | 7 |
|  |  | Rio Grande do Norte | 1 |
|  |  | Sergipe | 10 |
|  |  | Rio de Janeiro | 10 |
|  |  | Espírito Santo | 10 |
| **Hawksbill turtle**<br>(*Eretmochelys imbricata*) | Foraging Area | Fernando de Noronha | 7 |
|  |  | Atol das Rocas | 6 |
|  | Nesting Area | Bahia | 13 |
|  |  | Rio de Janeiro | 1 |
|  |  | Sergipe | 1 |
|  |  | Rio Grande do Norte | 11 |
| **Green turtle**<br>(*Chelonia mydas*) | Foraging Area | Fernando de Noronha | 1 |
|  |  | Ilha do Arvoredo | 1 |
| **Olive ridley turtle**<br>(*Lepidochelys olivacea*) | Nesting Area | Sergipe | 7 |
|  | Foraging Area | Sergipe | 2 |
| **Leatherback turtle**<br>(*Dermochelys coriacea*) | Foraging Area | Ceará | 1 |
|  |  | Pesca | 1 |
| **Hybrid *Cc* x *Ei*** | Nesting Area | Bahia | 17 |
|  | Foraging Area | Ceará | 2 |
|  |  | Atol das Rocas | 1 |
|  |  | Sergipe | 1 |
| **Hybrid *Ei* x *Lo*** | Nesting Area | Bahia | 2 |
| **Hybrid *Cc* x *Lo*** | Foraging Area | São Paulo | 1 |
|  | Nesting Area | Sergipe | 10 |
| **Hybrid *Ei* x *Cc* x *Cm*** | Nesting Area | Bahia | 4 |

### Discovery and standardization of nuclear markers

The initial selection of nDNA markers was performed using a reduced genomic dataset produced via ddRAD. As a preliminary analysis to help establishing a standardized ddRAD protocol for sea turtles (Driller et al. 2020), one individual of each parental species (*E. imbricata* and *C. caretta)* and one F1 *Ei* × *Cc* hybrid individual were used for ddRAD library construction. The sequencing library was generated by digesting the genomic DNA using the restriction enzymes *Nde*I and *Mlu*CI with subsequent ligation of Illumina adapters followed by a 10-cycle Polymerase Chain Reaction (PCR) for completeness of sequencing adapters, as described in Peterson et al. (2012). Libraries were pooled and size-selected between 500-600 bp using the PippinPrep equipment (Sage Science). Sequencing was performed on an Illumina MiSeq machine using a 600-cycle kit with the 300 bp paired-end sequencing mode. Samples were demultiplexed and the three samples were run through the pyRAD (Eaton 2014) pipeline for homologous loci recognition and genotyping. Briefly, the 300 bp paired-end reads were merged using PEAR (Zhang et al. 2013) and aligned into single sequences and those were clustered at 85% identity with a minimum coverage of ten and a maximum of five heterozygous sites per locus. The selected loci were manually screened for interspecific variation between the two species, and their sequences were extracted from the pyRAD output file and aligned using MUSCLE (Edgar 2004). For subsequent primer design, we selected only loci found in both parental species and the hybrid individual, showing a maximum of two indels and at least two interspecific differences between *E. imbricata* and *C. caretta*, which were confirmed as heterozygous in the hybrid. Primers were designed using the Primer3 (Untergasser et al. 2012) algorithm implemented in Geneious 8.1 (Kearse et al. 2012) using default parameters (Supplementary Table S2). Thus, we selected initially 24 anonymous nDNA markers with interspecific variation to be further validated for population studies.

The validation of ddRAD-derived nDNA markers was made through PCR amplification and Sanger sequencing. PCR was done in a final volume of 20 μl using 200 μM dNTP, 0.5 units of Platinum™ Taq DNA polymerase (Life Technologies™), 1.5 mM of MgCl_2_, 0.5 μM of forward and reverse primers and 10 ng of genomic DNA in 1X reaction buffer. PCR conditions were performed with one initial denaturation cycle of 95°C for 5 minutes, 35 cycles of denaturation of 95°C for 30 seconds, variable annealing temperatures (Supplementary Table S2) for 40 seconds, extension at 72 °C for 1 minute, and a final extension at 72 °C for 7 minutes. PCR products were purified by precipitation using a solution of 20 mM polyethylene glycol and 2.5 mM NaCl.

The Sanger sequencing reaction was performed using the BigDye Terminator Cycle Sequencing kit (Applied Biosystems™) following the manufacturer’s standard protocol. Forward and reverse sequences were generated on the ABI 3130xl DNA sequencer (Applied Biosystems™). The SeqScape v2.6 software (Applied Biosystems™) was used to check the electropherogram quality. Heterozygous sites were verified for accuracy and coded as ambiguous sites according to IUPAC code. High quality consensus sequences were aligned using the ClustalW algorithm in the MEGA 7 software (Kumar et al. 2016). The PHASE algorithm (Stephens et al. 2001) was used for gametic phase reconstruction of the heterozygous sequences with the assistance of Seq-PHASE input/output interconversion tool (Flot 2010). The DnaSP v5 program (Librado and Rozas 2009) was used for haplotype assignment. Heterozygous indels found in some sequences of locus 966 were phased using the Indelligent web tool (Dmitriev and Rakitov 2008). Finally, high quality phased sequences were verified again with the overlapping sequence chromatographs to edit for any inconsistencies.

Fifteen individuals of *E. imbricata*, 15 *C. caretta*, two *L. olivacea*, two green turtles (*Chelonia mydas*) and two leatherback turtles (*Dermochelys coriacea*), as well as ten hybrids, were initially sequenced for the 24 selected loci. Based on the intra and interspecific variation found, 15 out of the 24 loci were selected to be analyzed in a greater number of individuals (Supplementary Table S2). The most variable loci were selected for intraspecific analyses with the *C. caretta* (11 loci) and *E. imbricata* (14 loci) species (Table 2), while loci with greater power to distinguish different species were used for hybrid and phylogenetic analysis (Supplementary Table S2). Thus, each dataset was developed specifically for intra and interspecific studies of the target species using variation analyzed as high quality phased haplotypes obtained by Sanger resequencing, minimizing the ascertainment bias.

**Table 2.** Genetic diversity of 15 nuclear markers selected for *C. caretta* (11) and *E. imbricata* (14) intraspecific analyses. Number of individuals (N), number of haplotypes (H), number of polymorphic sites (S) and haplotype diversity (k). Uninformative marker (−).

|  |  | 421 | 856 | 966 | 3061 | 9672 | 23712 | 30573 | 31476 | 42006 | 46208 | 67959 | 76958 | 109472 | 114650 | 267557 |
| --- | --- | --- | --- | --- | --- | --- | --- | --- | --- | --- | --- | --- | --- | --- | --- | --- |
| *Eretmochelys imbricata* | N | 39 | 39 | 39 | 35 | - | 39 | 39 | 34 | 39 | 39 | 35 | 35 | 39 | 39 | 39 |
|  | H | 3 | 2 | 2 | 2 | - | 3 | 5 | 2 | 3 | 3 | 2 | 4 | 4 | 3 | 5 |
|  | S | 3 | 1 | 3 | 1 | - | 2 | 3 | 4 | 2 | 3 | 2 | 4 | 3 | 2 | 4 |
|  | k | 0.48 | 0.46 | 0.41 | 0.11 | - | 0.39 | 0.71 | 0.47 | 0.21 | 0.54 | 0.32 | 0.21 | 0.6 | 0.49 | 0.3 |
| *Caretta caretta* | N | 53 | 53 | 53 | - | 49 | 53 | 53 | - | 53 | 53 | - | - | 52 | 53 | 47 |
|  | H | 2 | 3 | 6 | - | 4 | 3 | 2 | - | 5 | 4 | - | - | 4 | 3 | 7 |
|  | S | 1 | 3 | 5 | - | 3 | 2 | 1 | - | 5 | 3 | - | - | 3 | 2 | 6 |
|  | k | 0.17 | 0.12 | 0.47 | - | 0.44 | 0.07 | 0.05 | - | 0.67 | 0.23 | - | - | 0.54 | 0.51 | 0.59 |

All sequences generated in this study have been deposited in GenBank and this research is registered in the National System for Genetic Heritage and Associated Traditional Knowledge (SisGen) of Brazil under number A03A2C2.

### Analyses of hybrids

A phylogenetic network was built to represent the interspecific lineage admixture of hybrid individuals (Joly et al. 2015). We estimated genetic distances (Joly et al. 2015) using a distance matrix of alleles and converting it into a distance matrix of individuals using the program POFAD (Joly and Bruneau 2006). We used the MEGA 7 software (Kumar et al. 2016) to generate genetic distances using Kimura-2-parameters model for each of the 14 loci (Dataset 1 in Supplementary Table S2) and then generated a combined-locus distance matrix using POFAD. The resulting matrix was used to build a phylogenetic network (neighborNet) using the software SplitsTree 4 (Huson and Bryant 2006).

Bayesian clustering analysis was done in the STRUCTURE software (Pritchard et al. 2000) for inference on population structure and assignment of individuals to populations using multilocus data. We assumed the admixture model where the individuals may have mixed ancestry in more than one of the K populations (species), allowing detection of the introgression level (Pritchard et al. 2000; Falush et al. 2003). Five loci were excluded from the STRUCTURE analysis because they present either a high-level of shared haplotypes between species or a considerable level of missing data. Two individuals of *D. coriacea* were also excluded due to a large amount of missing data, likely due to the low level of homology in the selected primers originally designed from *E. imbricata* and *C. caretta* sequences. The final dataset was composed by haplotypic data inferred for nine nDNA loci (Dataset 2 in Supplementary Table S2) genotyped in individuals from rookeries and feeding areas from four sea turtle species. We also performed intraspecific analyses using the datasets including 11 loci for *C. caretta* and 14 loci for *E. imbricata* (Table 2). Twenty independent runs for each K value (from K=1 to K=7) were performed with 200,000 Markov Chain Monte Carlo (MCMC) repeats after a 100,000 burn-in period. The independent and correlated allele frequencies were tested. The best K was assessed using Evanno’s methodology (Evanno et al. 2005) through the online tool STRUCTURE Harvester (Earl and VonHoldt 2012). We combined the replicate result files and visualized the estimated membership coefficients using CLUMPAK (Kopelman et al. 2015).

The posterior probability of each individual to belong to different hybrid classes was analyzed in NewHybrids v. 1.1 Beta3 (Anderson and Thompson 2002). Separate datasets combining different hybrid crossings were tested, since the NewHybrids only consider hybridization events involving two diploid species (Anderson 2008). Therefore, individuals resulted from crosses involving likely more than two species (R0264 and R0265) could not be analyzed. The analysis was done using the Jeffrey option, no priors, with a burn-in period of 100,000 and 500,000 MCMC sweeps. The following genotype classes were considered: pure parental (Pure 1 and Pure 2), first and second-generation hybrids (F1 and F2 between two F1 hybrids) and backcrosses between F1 and pure parental (BC1 and BC2). The R package HybridDetective was used to plot NewHybrids analysis (Wringe et al. 2017).

### Genetic diversity and population structure

Population analyses were performed for *C. caretta* and *E. imbricata* using nDNA markers with larger intraspecific variation. Diversity indexes were generated using the Arlequin v3.5 (Excoffier and Lischer 2010), DnaSP and MEGA software. The summary statistics used were: number of haplotypes (H), haplotype diversity (k) and number of polymorphic sites (S). Principal Component Analysis (PCA) was performed using R package *adegenet* to evaluate the genetic diversity among the sampled individuals (Jombart and Ahmed 2011). The missing data was replaced by the mean allele frequency and the PCA of standardized allele frequencies at the individual level was calculated using multivariate methods without spatial components. The analyses were performed including either (*i*) individuals collected in nesting and feeding areas along the Brazilian coast, or (*ii*) only females sampled in Brazilian nesting areas.

To investigate the relationship of different lineages and to represent part of the worldwide genetic diversity within species, mitochondrial control region haplotypes were compiled from literature and depicted in a network analysis. For *C. caretta*, the haplotypes (776 bp) were obtained from Shamblin et al. (2014), Nishizawa et al. (2014) and from the database of The Archie Carr Center for Sea Turtle Research (http://accstr.ufl.edu/resources/mtdna-sequences). For *E. imbricata*, the control region haplotypes (739 bp) were obtained from LeRoux et al. (2012), Vilaça et al. (2012), Vargas et al. (2016) and Gaos et al. (2018). The haplotype networks were constructed using the Reduced Median algorithm with reduction threshold 9 followed by Median Joining algorithm (RM-MJ network - Bandelt et al. 1995) using the software Network 5.0 (http://www.fluxus-engineering.com). The delimitation and nomenclature of the mtDNA clades were based on previous studies (LeRoux et al. 2012; Shamblin et al. 2014; Vargas et al. 2016) and are available in Supplementary Table S3.

To investigate the intraspecific multilocus allelic variation, we built a phylogenetic network from a combined-locus genetic distance matrix. We used the dataset of 11 loci for *C. caretta* and 14 for *E. imbricata*, and performed the network reconstruction using POFAD and SplitsTree 4 software as described above.

### Phylogenetic analysis

A phylogenetic reconstruction between sea turtle species was inferred using multilocus data with a Bayesian method implemented in BEAST v2.4.3 (Bouckaert et al. 2014). The sequences of 14 anonymous loci (Dataset 1 in Supplementary Table S2) were analyzed for five species of sea turtles.

The selection of partitioned models of molecular evolution was made using the PartitionFinder2 software (Lanfear et al. 2017). The best-fit model was selected by AICc criterion (Supplementary Table S4). The phylogenetic tree was inferred assuming a relaxed lognormal molecular clock under the birth-death model. This diversification model assumes that each species has a constant probability of speciating or going extinct along the lineage. It was employed considering that sequences from different species were used, and the species were sampled in different levels and presented very different branch length (Drummond and Bouckaert 2014). Fossil and genetic evidence (Bowen et al. 1993; Duchene et al. 2012) provided reference dates to be used as priors for tree calibration with a lognormal distribution, as follows: *i*) split between Dermochelidae and Cheloniidae family was set to 115 million years ago (mya) with a 95% confidence interval of 106–130 mya (Hirayama 1998) and *ii*) Carettini and Chelonini tribe was set to 65 mya with a 95% confidence interval of 50–90 mya (Moody 1974; Cadena and Parham 2015). The monophyly of the ingroup (Cheloniidae) was assumed a priori, by using *D. coriacea* as outgroup. The estimated date should be interpreted as maximum age constraints of the nodes.

Three independent MCMC chains were run for 200,000,000 generations and sampled every 5,000 generations. Trace files were checked for chain convergence and sufficient effective sample sizes (ESS) in Tracer v. 1.6 (http://beast.bio.ed.ac.uk/Tracer), considering ESS>200 as acceptable. The maximum clade credibility (MCC) tree was summarized after a 50% burn-in in TreeAnnotator from the 20,000 trees.

## Results

### Analyses of hybridization

High quality multilocus data standardized in this work was used to identify hybrids and estimate the introgression level in sea turtles. Some nDNA markers were more informative to characterize hybrids since they presented species-specific haplotypes that allowed us to identify the parental origin of the alleles with greater confidence. Loci 856, 3061, 76958 and 109472 were analyzed for a greater number of individuals and presented a larger number of diagnostic sites to identify *Cc* × *Ei* hybrids, while loci 421, 3061 and 109472 have greater power to identify *Lo* × *Ei* hybrids, and loci 421, 966, 67959 and 114650 to identify *Cc* × *Lo* hybrids (Supplementary Table S2).

Considering the combined data from all 14 nDNA loci selected with interspecific differences, the POFAD analysis produced a reticulated network of all individuals. The five sea turtle species were recovered in different clusters and the hybrids were observed in an intermediate position between species involved in the hybridization process (Figure 1). This method allowed the characterization of the genomic admixture of hybrids using distance measures to estimate the contribution of parental genomes.

**Figure 1.**
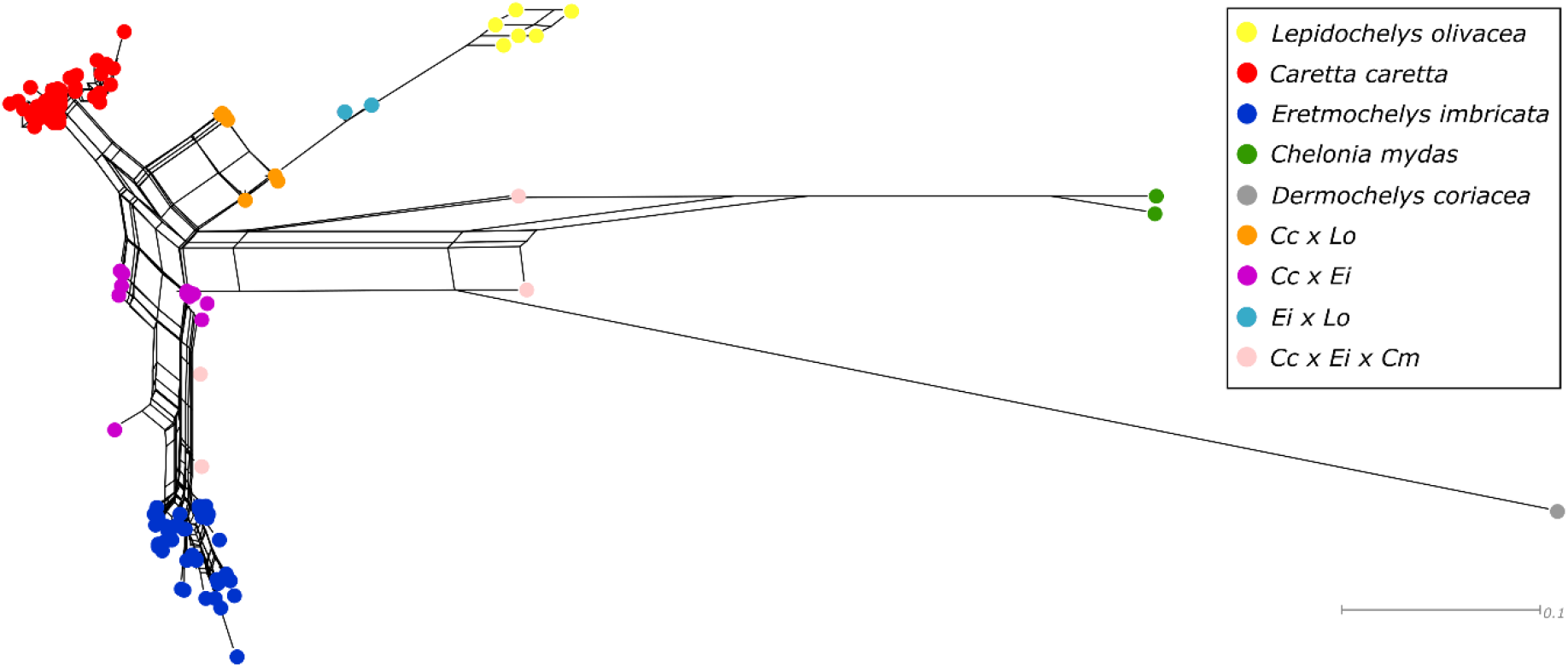
NeighborNet of organisms based on multilocus nuclear data for sea turtle species and hybrids that occur along the Brazilian coast. The hybrids are observed intermediately between species involved in the hybridization process. Details of sampling (N=143) are described in Table 1. Tips of the neighborNet represent unique multilocus genotypes. Cc: *Caretta caretta*, Ei: *Eretmochelys imbricata*, Lo: *Lepidochelys olivacea*, Cm: *Chelonia mydas*.

The Bayesian clustering analysis generated by STRUCTURE using correlated allele frequencies model showed that the number of clusters that best fit the data according to the Evanno’s statistics (Evanno et al. 2005) was five (Figure 2). The four sea turtle species included in this analysis were distinguished in different groups with high probability (99.9%) according to NewHybrids. *Caretta caretta* individuals were clustered in two different subgroups, one corresponding to individuals with mtDNA haplotypes commonly found in Brazilian rookeries, and another including foraging individuals sampled at Elevação do Rio Grande (ERG) with mtDNA haplotypes found in rookeries of the Caribbean, Mediterranean and Indo-Pacific oceans. The same intraspecific subdivision was obtained when *C. caretta* individuals were analyzed separately (Supplementary Figure S1), but it was not observed using independent allele frequency model (Supplementary Figure S2). All individuals of *E. imbricata* from Brazil were attributed to a single population in STRUCTURE analysis using both multi-species (Figure 2) and intraspecific (Supplementary Figure S1) datasets.

**Figure 2.**
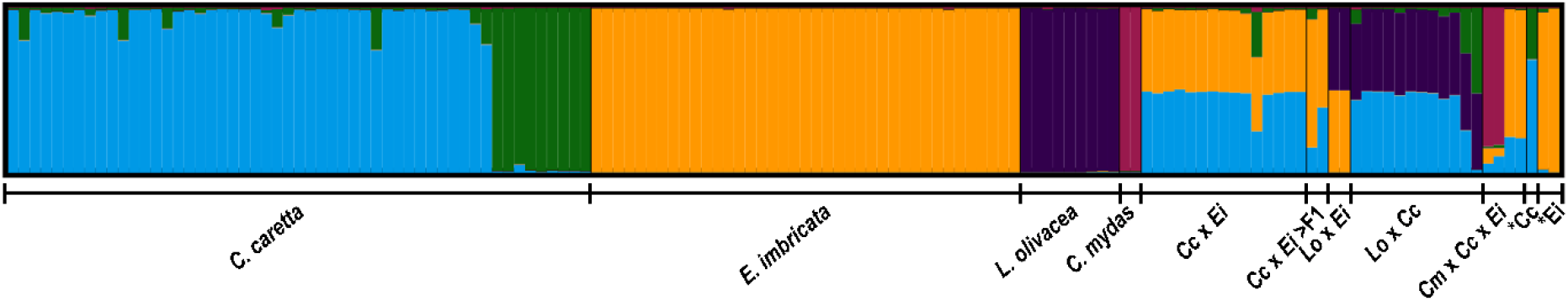
STRUCTURE bar plots representing K = 5 using correlated allele frequencies model using nine nuclear markers. The x-axis represents each individual analyzed and the y-axis represents the estimated admixture proportions related to each parental species. The barplot was obtained with CLUMPAK. The asterisks (*) show misidentified individuals. Cc: *Caretta caretta*, Ei: *Eretmochelys imbricata*, Lo: *Lepidochelys olivacea*, Cm: *Chelonia mydas*, >F1: introgressed hybrid.

Since the admixture model was assumed in STRUCTURE, the introgression level of hybrids could be also inferred. F1 hybrids clearly displayed intermediary genomic composition between parental species. All *Cc* × *Lo* and *Ei* × *Lo* hybrids were classified as F1 with a probability of 99.9% according to NewHybrids analysis. The *Cc* × *Ei* hybrids identified as F1 presented a posterior probability of 99.9% (NewHybrids) of belonging to this category. The parental *C. caretta* population involved in hybridization cases is associated with mtDNA haplotypes typically found in Brazil.

Three individuals (R0069, R0072 and R0217) previously assigned as hybrids (>F1, F1 and >F1, respectively) by Vilaça et al. (2012) showed no evidence of admixture between species for all nine nDNA markers analyzed. Individuals R0069 and R0072 were identified as *E. imbricata* and R0217 as *C. caretta* with high posterior probability. This could have resulted from sample misidentification, as we confirmed by re-sequencing the loci RAG1 and CMOS used by Vilaça et al. (2012), which reinforced that they are indeed ‘pure’ individuals (Supplementary Table S5). We have also re-sequenced the control region of mtDNA for individual R0072 and, in contrast to the previous work, it presents a haplotype from *E. imbricata*. For the individual R0217 that was “morphologically” identified as *E. imbricata*, all the genetic data suggest that it is a pure *C. caretta*, probably due to misidentification during subsequent sample manipulation.

Previous work (Vilaça et al. 2012) identified 17 individuals as introgressed (>F1) hybrids, of which 15 were re-analyzed in this work with a multilocus nDNA approach. Using our nDNA dataset, we were able to recognize only six individuals with evidence of being >F1 generation hybrids (Supplementary Figure S3 and Table S1). Remarkably, they were all hatchlings collected in nests and showing characteristics of more than one sea turtle species (Vilaça et al. 2012). Individual R0025 is a hatchling of a *Cc* × *Ei* hybrid female (R0024) and it was attributed to the category of backcrossing with an *E. imbricata* male with a probability of 99.8% (NewHybrids). Individual R0196 was sampled with morphological evidence of hybridization and it was also identified as backcrossing with an *E. imbricata* male with a probability of 96.8% (NewHybrids). The remaining four hatchlings (R0264, R0265, R0267 and R0268) are siblings derived from a single clutch. The genetic admixture of three species *E. imbricata* × *C. caretta* × *C. mydas* (*Ei* × *Cc* × *Cm*) was confirmed in two individuals (R0264 and R0265), although the posterior probability could not be estimated because NewHybrids only considers hybridization cases involving two species. The remaining siblings R0267 and R0268 were attributed to the category backcrossing with *E. imbricata* with a posterior probability of 99.8% (NewHybrids). This result is in accordance with Vilaça et al. (2012) which hypothesized that these hatchlings could have resulted from the crossing between one *Cc* × *Ei* F1 hybrid female with at least one *C. mydas* male (evidenced by the R0264 and R0265) and another *E. imbricata* male (evidenced by the R0267 and R0268).

### Population analyses

Population analyses were performed independently for *C. caretta* and *E. imbricata* using the intraspecific nuclear variation. A total of 4492 bp were sequenced from 14 nDNA markers for *E. imbricata* and 3592 bp were sequenced from 11 nDNA markers for *C. caretta* (Table 2).

A PCA of multilocus data (Figure 3) was conducted to infer population structure assessing continuous axes of genetic variation of these species. First, we investigated nesting areas along the Brazilian coast for *C. caretta*. The first axis explained only 33.8% of the total variation, while the second axis explained 29.8% of the variation, showing no relevant structure between Brazilian populations (Figure 3C). When individuals sampled in feeding areas were included in the analysis, the first principal component (PC1) explained 69.7% of the total variation and divided the samples into two clusters (Figure 3D). The first one corresponds to all individuals from Brazilian rookeries and nine individuals captured in the feeding area in southern Brazil – Elevação do Rio Grande (ERG) – that present mtDNA haplotypes commonly found in Brazilian rookeries (CC-A4 and CC-A24). The second one corresponds to individuals from ERG that present mitochondrial haplotypes (CC-A11, CC-A2, CC-A33 and CC-A34) found in rookeries of the Caribbean, Mediterranean and Indo-Pacific oceans. The second principal component (PC2) represents 29.7% of the variation.

**Figure 3.**
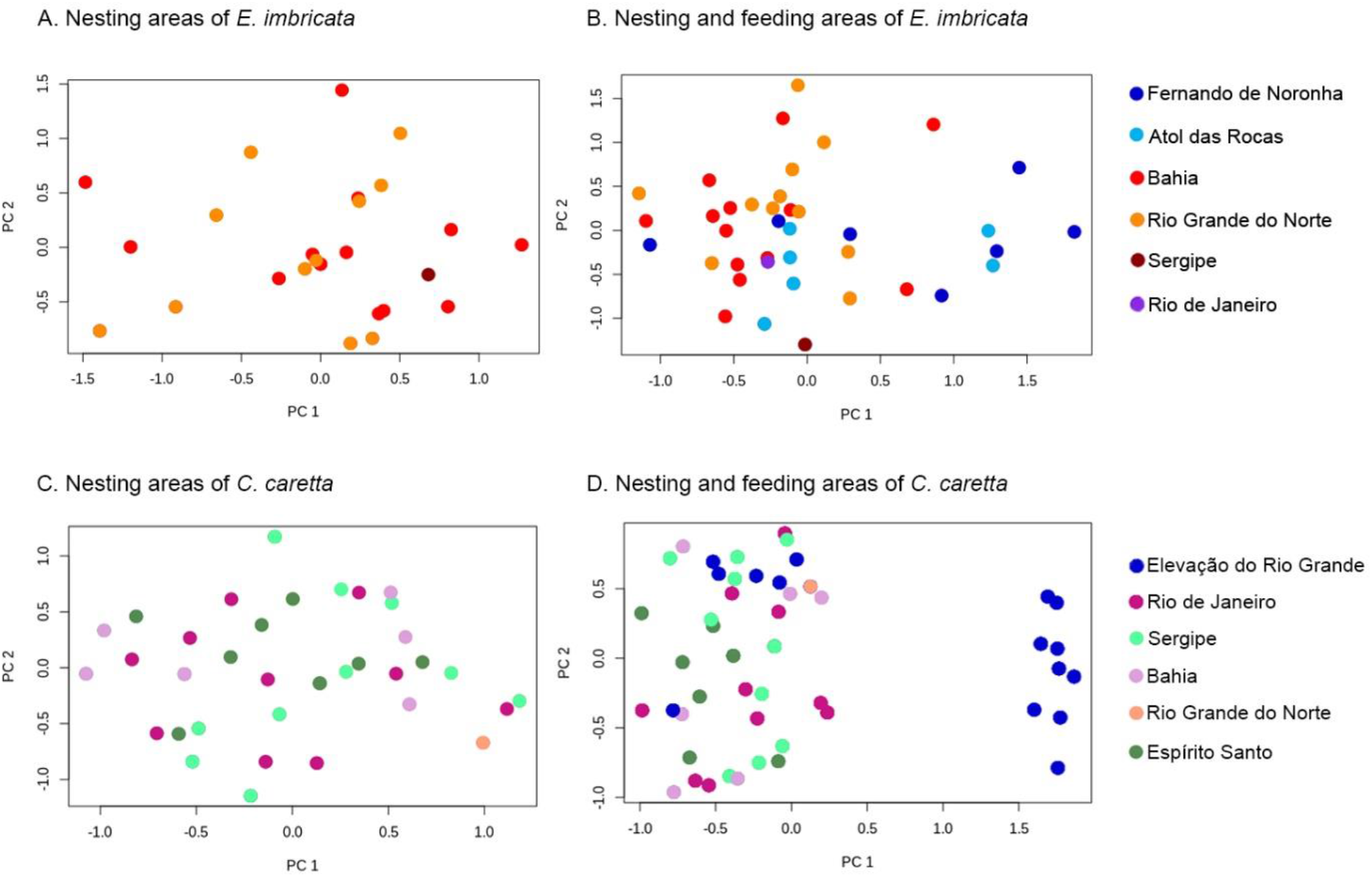
Principal component analysis of multilocus data for *C. caretta* (11 markers) and *E. imbricata* (14 markers) including only sea turtles sampled in Brazilian rookeries (A and C) and individuals collected in rookeries and feeding areas along the Brazilian coast (B and D). Color codes indicate the geographical location where the individuals were collected.

A PCA was performed for individuals of *E. imbricata* sampled in Brazilian rookeries. PC1 and PC2 explained 46.12% and 34.87% of the total variation, showing no correlation between genetic variation and geographic distribution (Figure 3A). When sea turtles from feeding areas were included, PC1 and PC2 explained 49.77% and 40.44% of the total variation, respectively (Figure 3B). Five individuals sampled at feeding areas of Fernando de Noronha and Atol das Rocas were slightly separated from other individuals. They presented mtDNA haplotypes either typically found in the Indo-Pacific Ocean basin (EiIP16 and EiIP33) or ‘orphan’ haplotypes (EiA49, EiA75) which are differentiated from EiIP16 by one mutation step.

Population structure was analyzed comparing mtDNA and nDNA data of *C. caretta* (N=53) and *E. imbricata* (N=39) from the Brazilian coast. The 98 mtDNA haplotypes of *C. caretta* compiled from the literature were depicted in the network (Figure 4A), which showed three clades representing the main lineages within species (Shamblin et al. 2014). There was a large genetic divergence between mtDNA clades, a pattern not observed with our multilocus data (Figure 4B). The neighborNet showed that some individuals from different mtDNA clades exhibited a close phylogenetic relationship when nDNA data was considered (highlighted with an ellipse in Figure 4B).

**Figure 4.**
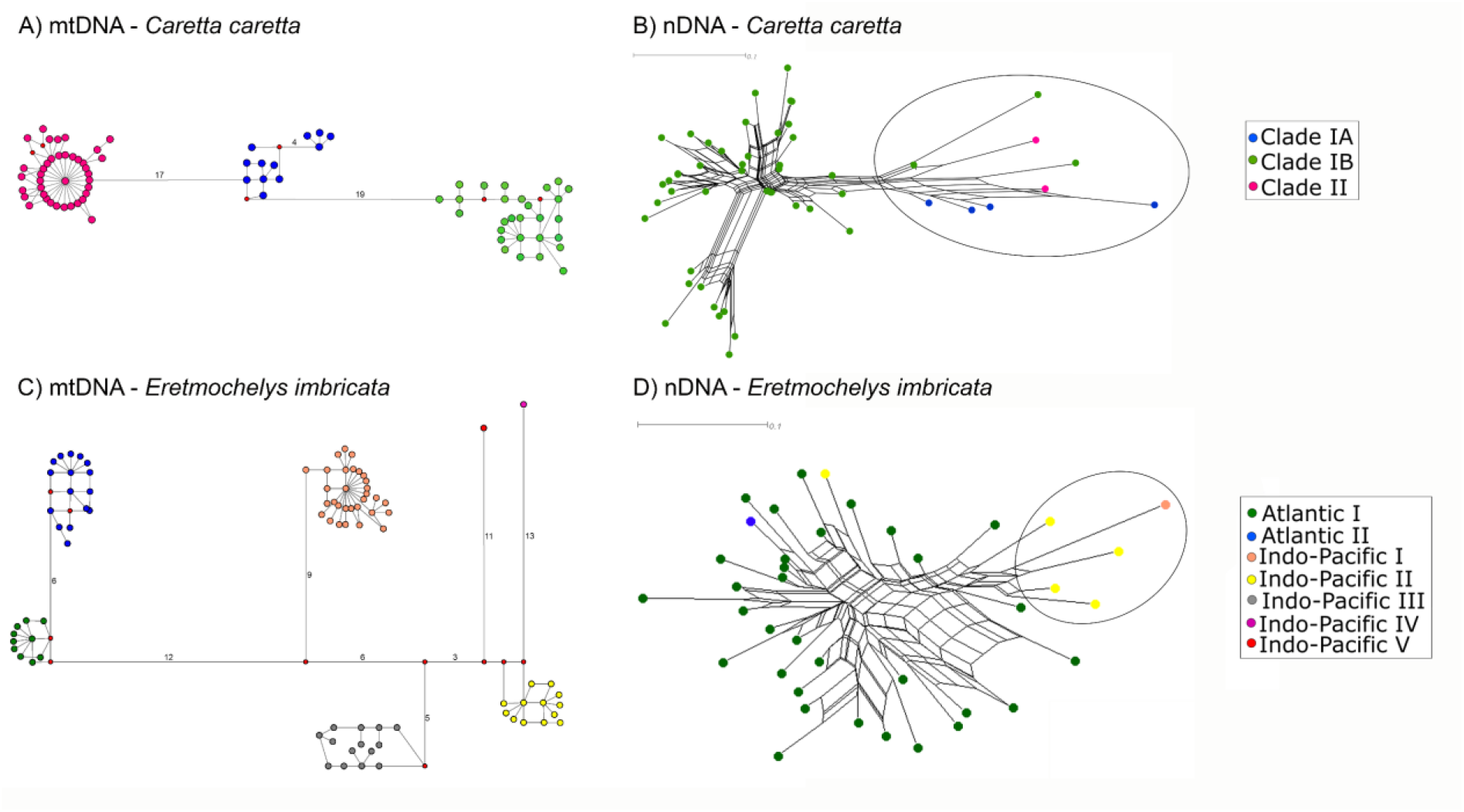
Haplotype network based on mtDNA control region data (A and C) and neighborNet of organisms based on multilocus nuclear data (B and D) for *C. caretta* and *E. imbricata*. The mitochondrial data were obtained from haplotypes based on control region previously published in literature. The nuclear data comprised 11 loci for 53 individuals of *C. caretta* and 14 loci for 39 individuals of *E. imbricata*. Tips of the neighborNet represent unique multilocus genotypes. The ellipses highlight the individuals of *C. caretta* more distantly related and supposed to have Indo-Pacific origin (B) and the individuals of *E. imbricata* that belong to the Indo-Pacific mtDNA clades and were grouped together (D).

For *E. imbricata*, the relationship among 87 control region mtDNA haplotypes obtained from literature was depicted in a network shown in Figure 4C. They were clustered in seven main clades, two reported in the Atlantic Ocean and five in the Indo-Pacific Ocean. The neighborNet built with multilocus data did not present large genetic distances between individuals from Atlantic and Indo-Pacific (Figure 4D). However, five of six individuals collected in foraging aggregations in northern Brazilian coast that belong to Indo-Pacific mtDNA clades were clustered in an end of the neighborNet (highlighted with an ellipse in Figure 4D), suggesting they come from another gene pool. Only individual R0242 from the Indo-Pacific mtDNA clade appeared more closely related to individuals that belong to the Atlantic mtDNA clade.

### Phylogenetic analysis

The MCC tree obtained with multilocus data (Figure 5) showed the topology and dating congruent with previous phylogenetic studies of sea turtles (Bowen et al. 1993; Naro-Maciel et al. 2008; Duchene et al. 2012). The estimation of the time to most recent common ancestors (TMRCA) for the five species of sea turtles was 112.4 mya. The divergence between Carettini and Chelonini tribe (Cheloniidae) was estimated to have occurred at 65.9 mya. *Eretmochelys imbricata* separated from *C. caretta* and *L. olivacea* at 25 mya, followed by the split between *C. caretta* and *L. olivacea* at 21.6 mya.

**Figure 5.**
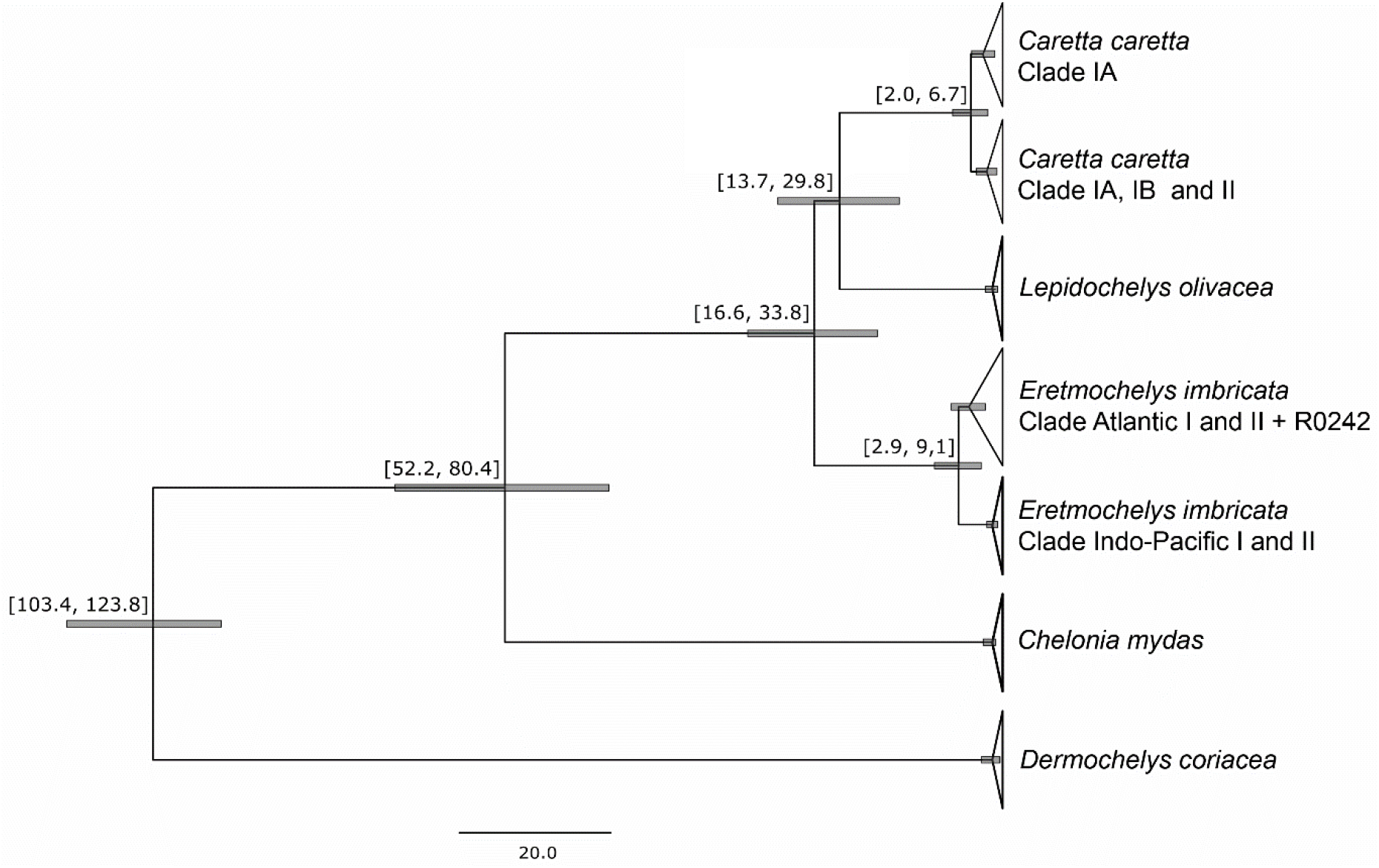
Dated Bayesian phylogeny of sea turtles from the Brazilian coast inferred from multilocus data. The horizontal axis indicates divergence times in million years before present. Horizontal bars and the numbers above branches correspond to the 95% highest posterior density (HPD) interval values estimated for all tree nodes with posterior probabilities above 0.8 calculated in BEAST. Clade names are based on mtDNA haplotypes as grouped by previous studies (LeRoux et al. 2012; Nishizawa et al. 2014; Shamblin et al. 2014; Vargas et al. 2016).

The divergence between Atlantic and Indo-Pacific lineages of *E. imbricata* was estimated to have occurred at 5.93 mya, approximately the same date estimated using control region haplotypes (Vargas et al. 2016). Monophyly of clades based on mtDNA of *E. imbricata* (LeRoux et al. 2012; Vargas et al. 2016) was supported with nDNA in the Bayesian analysis using BEAST, except for one individual (R0242). This sea turtle belongs to mtDNA Indo-Pacific clade II, but its nuclear composition showed that it is more similar to individuals from the Atlantic mtDNA clade. However, R0242 represents an early nDNA diverging branch in the Atlantic mtDNA clade, despite the low clade Bayesian posterior probability (0.41).

The earliest divergence between *C. caretta* lineages was estimated to have occurred 4.29 mya, similar to the date estimated using mitogenomes (Duchene et al. 2012) and control region haplotypes (Shamblin et al. 2014). One nDNA lineage gathers nesting and foraging individuals from Brazil that presents mtDNA haplotypes derived from CC-A4 and the other nDNA lineage presents individuals foraging in ERG that belong to three different mtDNA clades (IA, IB and II). Thus, the mtDNA based clades were only partially recovered with nuclear multilocus data, since individuals from different mtDNA clades were grouped in a single nDNA lineage.

## Discussion

### The interspecific hybridization phenomenon along the Brazilian coast

The use of a multilocus approach resulted in an informative dataset based on high quality haplotypes that allowed expanding our comprehension about the hybridization process of sea turtles. In this study, we re-analyzed 15 out of 17 individuals previously identified as introgressed (>F1) hybrids by Vilaça et al. (2012), but we only confirmed six backcrossed individuals. All the introgressed (F2 hybrids) individuals were hatchlings. Seven F1 hybrids detected with our multilocus data were previously identified as introgressed (>F1) based on the information of only one genetic marker (Supplementary Table S1). It means that for all other 11 genetic markers analyzed by Vilaça et al. (2012), the individuals presented one allele of each parental species. For three individuals, the introgression signal was detected with one microsatellite or RFLP marker, which is based on allele size differences and present a high level of homoplasy and many genotyping artifacts such as null alleles and allele dropouts (Zhang and Hewitt 2003). Here we used nDNA multilocus resequencing to characterize high quality haplotypes that supply a much higher level of resolution at both inter and intraspecific analyses (Schlötterer 2004). The use of genetic markers randomly distributed throughout the genome, generated by high quality Sanger sequencing data, provided the highest genotyping accuracy with low ascertainment bias. Indeed, our multilocus dataset displayed a higher power to distinguish different hybrid crossings and introgression levels as compared to the previous methods.

Even though the initial NGS screening of variable loci was done using the ddRAD approach with only two species (*C. caretta* and *E. imbricata*), we were able to validate informative nDNA loci with diagnostic alleles/haplotypes for other Chelonioidea taxa, even for species displaying close phylogenetic relationship as *L. olivacea* and *C. caretta*. Considering the number of individuals analyzed and the number of diagnostic sites, we suggest the use of loci 856, 3061, 76958 and 109472 to characterize *Cc* × *Ei* hybrids, loci 421, 3061 and 109472 to characterize *Ei* × *Lo* hybrids, and loci 421, 966, 67959 and 114650 to characterize *Cc* × *Lo* hybrids (Supplementary Table S2). Future genetic studies investigating the hybridization between different species of sea turtles should be able to select more informative loci according to their target species.

According to Vilaça et al. (2012), there are introgressed (at least F2 hybrids) adult females nesting in Bahia (Brazil), and the first interspecific crossing could have occurred at two generations ago or a minimum of 40 years. In contrast, our data suggest that only hatchlings (newborns) were confirmed as introgressed hybrids. Considering the age at maturity from 20 to 40 years for *E. imbricata* (Meylan and Donnelly 1999) and from 22 to 29 years for *C. caretta* (Heppel 1998; Casale et al. 2011), we estimate that the minimum time for the first hybridization event was one generation ago (at least 20 years). Since the first hybrid female analyzed in this work was sampled in year 2000 at Bahia, our data suggest that the high-frequency hybridization event in Bahia may have started around 1980. This is supported by Conceição et al. (1990), which in 1989 first recorded hybrid juveniles in the state of Bahia. Bass et al. (1996) also support this hypothesis since they were the first genetic work to report, in 1992, a high incidence of *Cc* × *Ei* hybrid hatchlings (10 of 14 individuals) of females morphologically identified as *E. imbricata* at Praia do Forte, Bahia. This indicates that introgression was likely overestimated by Vilaça et al. (2012) and hybridization may be a more recent phenomenon happening in Brazil. However, another possible hypothesis is that the hybridization may be a recurrent event, and the introgressed hybrids (F2) are much less fertile or inviable, precluding their survival and reproduction.

Studies have reported that the emergence success of hybrids is significantly lower than either hawksbills or loggerheads, although the hatchling production per clutch, breeding and nesting frequency, and hatchling viability of hybrids were similar to parental species (Soares et al. 2017; Soares et al. 2018). However, they only investigated F1 hybrids and their hatchlings. There is no information about the potential effects of hybridization in other life stages at the sea, as survivorship, growth rates and mating success. Indeed, if all (or the large majority) hybrid adults are first-generation hybrids as our results indicate, thus a most likely explanation is that outbreeding depression (decrease of survival and/or reproductive fitness) may occur mostly in the second and further generations of introgressed individuals. In this situation, the original parental gene combinations are broken up by recombination in >F1 hybrids, disrupting coadapted gene complex (Edmands et al. 1999; Goldberg et al. 2005).

The emergence of high-frequency hybridization cases in Brazil coincides with the period of a great population decline of sea turtles during the XX century. This depletion leads to a reduced chance of potential conspecific encounters, which may be associated with this unique event on the Brazilian coast (Vilaça et al. 2012). Reports of hybridization cases associated with human impact are increasing worldwide for other species (Allendorf et al. 2001; Grabenstein and Taylor 2018). Human activities may lead to secondary contact between previously isolated populations due to habitat disturbance and environmental changes that increase the hybrids rate (Todesco et al. 2016). Since 1980, sea turtle conservation in Brazil mostly relies on efforts of Projeto TAMAR, a consolidated and successful program aiming at environmental education and monitoring and research of sea turtles. Thereafter, the number of nesting females in monitored beaches has been increasing quickly (Marcovaldi and Chaloupka 2007), but in spite of this greater number of individuals, more recent hybridization events have been reported. A study of 2012 and 2013 nesting seasons showed that the incidence of hybridization in Bahia inferred from hatchlings of *C. caretta* females is 16.66% and for *E. imbricata* females is 8.15% (Soares et al. 2018).

Hybridization in Brazil is a local event with reports of fertile female hybrids in about 300 km of coastline between northern Bahia and Sergipe states. In this work, all female hybrids were originally sampled in rookeries of Bahia and Sergipe beaches, and pelagic individuals were sampled in coastal waters of Ceará, Bahia, Sergipe and São Paulo states. Other reports of hybrids in Brazil are juveniles from the states of Ceará and Rio Grande do Sul (Cassino Beach), which are two important feeding aggregations of *C. caretta* (Proietti et al. 2014). Further studies focusing on detailed characterization of hybrids is recommended, mainly in nesting areas worldwide with the overlapping distribution of different sea turtle species.

We confirmed that all the *Cc* × *Ei* F1 hybrids resulted from the crossing between *C. caretta* female and *E. imbricata* male, which indicates a gender bias. This is probably associated with the prevalence of *C. caretta* along the Brazilian coast and the partial overlapping of reproductive season with *E. imbricata* (Vilaça et al. 2012). The beginning of the nesting season for *E. imbricata* overlaps with the nesting peak of *C. caretta* (November and December), when *E. imbricata* males encounter a higher number of *C. caretta* females to mate (Proietti et al. 2014). Conversely, the encounter between *C. caretta* males and *E. imbricata* females may happen less frequently, since *C. caretta* males leave the mating areas before a large number of *E. imbricata* females arrive at nesting beaches (Vilaça et al. 2012).

Sea turtles present long and complex life cycles and monitoring the consequences of hybridization can be complicated, but it is extremely important to understand their impact on the management of sea turtle populations, particularly for parental species. Particular focus should be directed towards nests and hatchlings of F1 female hybrids to allow monitoring the future consequences of hybridization. We have shown that increasing the resolution of genetic data allows to better understand this local and atypical phenomenon in Brazil. New detailed genomic approaches should also be able to elucidate the relation between introgression and species-specific adaptive regions of the genome, in relation to lifecycle, foraging habitat and behavior of hybrids. Thus, further studies should be highly stimulated to expand our comprehension of this particular evolutionary process of potential conservation impact.

### Intraspecific studies of *C. caretta* and *E. imbricata*

Sea turtle genomic structure is quite monotonous, presenting slow cytogenetic and molecular divergence among species (FitzSimmons et al. 1995; Naro-Maciel et al. 2008; Wang et al. 2013). Indeed, our initial surveys of intra and interspecific variation for different sea turtles (Vilaça et al. 2012) revealed few informative nDNA markers. In this study, we were able to identify informative markers for intraspecific analyses that were validated after Sanger resequencing in some individuals of each sea turtle species. The nDNA intraspecific variation found in *C. caretta* and *E. imbricata* analyzed here (Table 2) allowed us to infer important patterns on population structure.

Unlike mtDNA, the nuclear loci isolated in this study showed that variation within both species is not significantly correlated to the geographic distribution along the Brazilian coast (Figure 3A and 3C, Supplementary Figure S1). Previous studies using mtDNA data showed significant differences in allelic frequencies between southern and northern Brazilian rookeries for *C. caretta* (Reis et al. 2010a; Shamblin et al. 2014) and *E. imbricata* (Vilaça et al. 2013). For *C. caretta*, three genetically distinct clusters based on mtDNA were recognized along the Brazilian coast: northern coast (Bahia and Sergipe), Espírito Santo and Rio de Janeiro (Shamblin et al. 2014). For *E. imbricata*, two different mtDNA clusters, although closely related, were reported: Bahia and Rio Grande do Norte (Vilaça et al. 2013).

Some discrepant results between mtDNA and nDNA analyses are expected, as these markers have different mutation rates, inheritance patterns and effective sizes. Because mtDNA follows a maternal inheritance, exhibits faster evolution rate and displays ¼ of the effective population size when compared with nDNA, it is used to investigate more recent demographic events (Cabanne et al. 2008; Brito and Edwards 2009). In contrast, nDNA reveals aspects from the biparental ancestry and a more ancient population history. Assessing the genetic diversity of nuclear markers is important for understanding the contribution of females and males to the population structure of sea turtles, as both sexes can have different behavior. However, regarding the definition of management units for conservation concerns, mtDNA data should be primordially considered since it characterizes relatively independent rookeries established by female philopatric recruitment (Shamblin et al. 2014).

Our population structure results can reflect an important sea turtle behavior. Lower population structure found in nDNA relative to mtDNA has been previously attributed to male-mediated gene flow (Bowen et al. 2005). Male sea turtles have uncertain philopatry and probably display greater flexibility in their choice of mating areas (FitzSimmons et al. 1997). Similar patterns were found in previous studies using nDNA and are indicative of lack of male philopatry (FitzSimmons et al. 1997; Bowen et al. 2005; Carreras et al. 2011; Vilaça et al. 2013; Clusa et al 2018). Thus, the apparently discrepant results for mtDNA and multilocus data could be further explained by male-mediated gene flow between rookeries. Future studies using genome-wide data should validate this hypothesis.

Considering feeding areas, the analysis of pelagic individuals was important to provide a better understanding of the genetic diversity of turtles, and to investigate migration routes and the connectivity of distant rookeries. Elevação do Rio Grande (ERG) area is a chain of undersea mountains located offshore of Brazil’s southern coast and is an important feeding aggregation and oceanic developmental habitat for immature *C. caretta*, probably attracted by the abundance of food. Confirming previous studies (Reis et al. 2010a; Shamblin et al. 2014), this area seems to be visited by individuals from nesting sites in Brazil, northern Atlantic Ocean and Indo-Pacific Ocean, which is an evidence for transoceanic migrations for the species via southern Africa. Origin inference of the pelagic individuals was made based on the mtDNA information, once individuals present typical haplotypes of specific rookeries. Two clusters were identified in the PCA of multilocus data, separating individuals of Brazilian rookeries, which present an exclusive mtDNA haplogroup CC-A4, from individuals originated in other continental rookeries (Figure 3C). The same clustering was observed for *C. caretta* in STRUCTURE analysis when correlated allele frequencies were used (Figure 2 and Supplementary Figure S1). In contrast, using independent allele frequencies model, this separation was not observed (Supplementary Figure S2). The correlated frequencies model provides greater power to identify distinct but closely related populations with recent shared ancestry (Porras-Hurtado et al. 2013).

For *E. imbricata*, all individuals were attributed to a single panmictic population in STRUCTURE analysis (Figure 2), even when it was performed including only *E. imbricata* individuals or using correlated allele frequencies model (Supplementary Figure S1). However, some individuals from feeding areas in northern Brazil exhibited mtDNA typically found in Indo-Pacific rookeries and were slightly separated in PCA (Figure 3D). The presence of individuals from distant rookeries, indicated in this work by nDNA, supports the occurrence of transoceanic migrations for this species.

Population structure was analyzed comparing mtDNA and nDNA data. Considering great part of mtDNA haplotype diversity reported in the literature for *C. caretta*, it is possible to distinguish three clades with great genetic divergence. They correspond to two major lineages - clades I and II - of which the former passed by a more recent split (subclades IA and IB, Figure 4A). In this work, subclade IA is represented by individuals with haplotypes CC-A33 and CC-A34, which were registered in Australian rookeries. Subclade IB is represented by individuals with mtDNA haplotypes derived from CC-A4, which were recorded only in Brazilian rookeries; CC-A1.3, reported in Florida-USA, Mexico and Cape Verde; and CC-A11.6, reported in Oman (Indian Ocean) (Shamblin et al. 2014). Clade II is represented by haplotype CC-A2.1, recorded in rookeries from South Africa, northwest Atlantic Ocean and Mediterranean Sea. Considering multilocus data, neighborNet analysis showed that nine individuals of *C. caretta* presented greater genetic divergence in relation to Brazilian individuals (highlighted with an ellipse in Figure 4B). These individuals were collected in the southern Brazilian feeding area (ERG) and belong to three different mtDNA clades (clades IA, IB and II). They also appear separately clustered in PCA (Figure 3D) and STRUCTURE (Supplementary Figure S1). Phylogenetic analysis also resulted in a MCC tree with two main lineages, of which one corresponds to a mix of individuals from three mtDNA clades (Figure 5).

Phylogeographic studies suggested that the two main mtDNA clades I and II of *C. caretta* were isolated by geographic and climatic factors into Atlantic and Indo-Pacific basins during the cooler periods of the Pleistocene (Bowen et al. 1994). As *C. caretta* also tolerates temperate water, migrations via southern Africa, directed by the waters of the Agulhas Current, are possible. The phylogeographic scenario proposed for Shamblin et al. (2014) suggests that mtDNA clade IA had an Indo-Pacific origin, where the earliest diverging lineages of *C. caretta* appear. The earliest colonization was likely from Indo-Pacific lineages invading the Atlantic Ocean. Brazilian haplotypes (CC-A4 and derived ones) seem to be the earliest diverging lineage within mtDNA clade IB, which was followed by a more recent colonization of the CC-A11.6 precursor from Atlantic to Indian Ocean, as it is closely related to Atlantic lineages. Therefore, transoceanic migration in both directions may be responsible for long-distance gene flow between *C. caretta* populations. Furthermore, current geographic distribution of these lineages presents no phylogenetic concordance, as both lineages are found in both Atlantic-Mediterranean and Indo-Pacific basins (Reis et al. 2010a; Duchene et al. 2012). Despite the small number of samples, this result can suggest a homogenization of *C. caretta* populations at a nuclear level for individuals sharing a common feeding area. This was previously reported for a *C. caretta* population in the southeastern USA and attributed to male-mediated gene flow (Bowen et al. 2005). This behavior should be elucidated using more representative data of the genetic variation through genomic surveys.

For *E. imbricata*, the relationship among previously reported mtDNA haplotypes revealed that there are seven main clades worldwide. Two of them were registered in rookeries from the Atlantic Ocean (LeRoux et al. 2012) and five in rookeries from the Indo-Pacific Ocean (Vargas et al. 2016). The neighborNet of nDNA data showed that 5 of 6 individuals of Indo-Pacific mtDNA clades are slightly more distant from individuals that belong to Atlantic mtDNA clades (Figure 4D). The same individuals belong to Indo-Pacific nDNA cluster according to the phylogenetic analyses (Figure 5). They were sampled in the Brazilian feeding aggregations, demonstrating long distance migrations for the species.

Despite the separation between *E. imbricata* individuals from Indo-Pacific and Atlantic was not clearly observed in STRUCTURE using nDNA, it was slightly observed in PCA and neighborNet, and strongly detected in the MCC tree. There is a deep genetic divergence between Indo-Pacific and Atlantic mtDNA lineages of *E. imbricata* dating from the early Pliocene, when the closing of the Isthmus of Panama occurs (Arantes et al. 2020). The geographic pattern of separation between ocean basins found with mtDNA was recovered with nuclear data, except for one individual (R0242). However, R0242 belongs to an early diverging lineage of the Atlantic mtDNA clade, which displays a low Bayesian posterior probability (0.41). It suggests that this individual is deeply related to all other mtDNA Atlantic lineages, thus a likely remnant of the first Indo-Pacific lineages colonizing the Atlantic Ocean.

*Eretmochelys imbricata* is strictly adapted to tropical waters and although some transoceanic migrations may occur, American and African continents are supposedly important barriers to species migration directing the current distribution of main lineages of sea turtles (Duchene et al. 2012). In contrast, *C. caretta* individuals were more divergent within the Atlantic than between the Atlantic and Indo-Pacific, probably due to a transoceanic gene flow observed in this species more adapted to temperate water. The barriers to gene flow are not the same for all species of sea turtles likely due to their different ability of dispersion through the oceans and evolutionary responses to environmental changes (Duchene et al. 2012).

Regarding intraspecific phylogenetic analysis, the use of multilocus data resulted in similar topology and divergence times between species when compared to the previous studies that used mtDNA data (Duchene et al. 2012; LeRoux et al. 2012; Shamblin et al. 2014; Vargas et al. 2016). The divergences between main lineages within *E. imbricata* and *C. caretta* were estimated to have occurred about 5.93 mya and 4.29 mya, respectively. It is consistent with the age of formation of the Isthmus of Panama, associated to the deepest phylogenetic split of intraspecific lineages of different species of sea turtles (Naro-Maciel et al. 2008; Duchene et al. 2012). Besides, assessing multilocus data made possible to evaluate biparental ancestry and accommodate the stochasticity of the coalescent process by combining information from multiple loci distributed throughout the genome, instead of relying only on inferences based on individual tree topologies (Edwards and Beerli 2000; Brito and Edwards 2009).

### Concluding remarks

Next Generation Sequencing technologies allowed the initial identification of genome-wide polymorphic loci, which were selected for a Sanger sequencing validation step to characterize multilocus datasets useful for inter and/or intraspecific studies. The high quality multilocus data provided significant interspecific information for the inference of the phylogeny of sea turtles and characterization of hybrids. Additionally, another multilocus dataset provided relevant intraspecific data for analyses of population structure. The presented results reveal important enhancements in the genetic resolution of the hybridization process and population structure of sea turtles.

## Acknowledgments

We are grateful for the access of collection samples from Projeto TAMAR, a conservation program of the Brazilian Ministry of the Environment, affiliated to ICMBio (the Chico Mendes Institute for Conservation of Biodiversity), and is co-managed by Fundação Pró-TAMAR and sponsored by Petrobras-Brazil. We are grateful for all members of Fundação Pró-TAMAR, particularly to M.A. Marcovaldi and L.S. Soares for their logistical support and availability of samples. We also thank Gisele Lobo-Hajdu from UERJ (Universidade Estadual do Rio de Janeiro) and Sarah M. Vargas (Universidade Federal do Espírito Santos) for sharing of previously collected samples. We also thank Christina Brown for helping with experiments and primer design. There were no new samples collected for this particular study.

## Data Accessibility

We have deposited the primary data underlying these analyses as follows:

- Genbank accession number for DNA sequences: MT23095-MT231002 and MT235978-MT23615.
- Sampling locations and individual data is available in Supplementary Tables S1 and S3.

