## Supplementary Figure S1 for "New genetic insights about hybridization and population structure of hawksbill and loggerhead turtles from Brazil"

**Arantes_SupMat_FigureS1_Hybridization of Brazilian sea turtles**

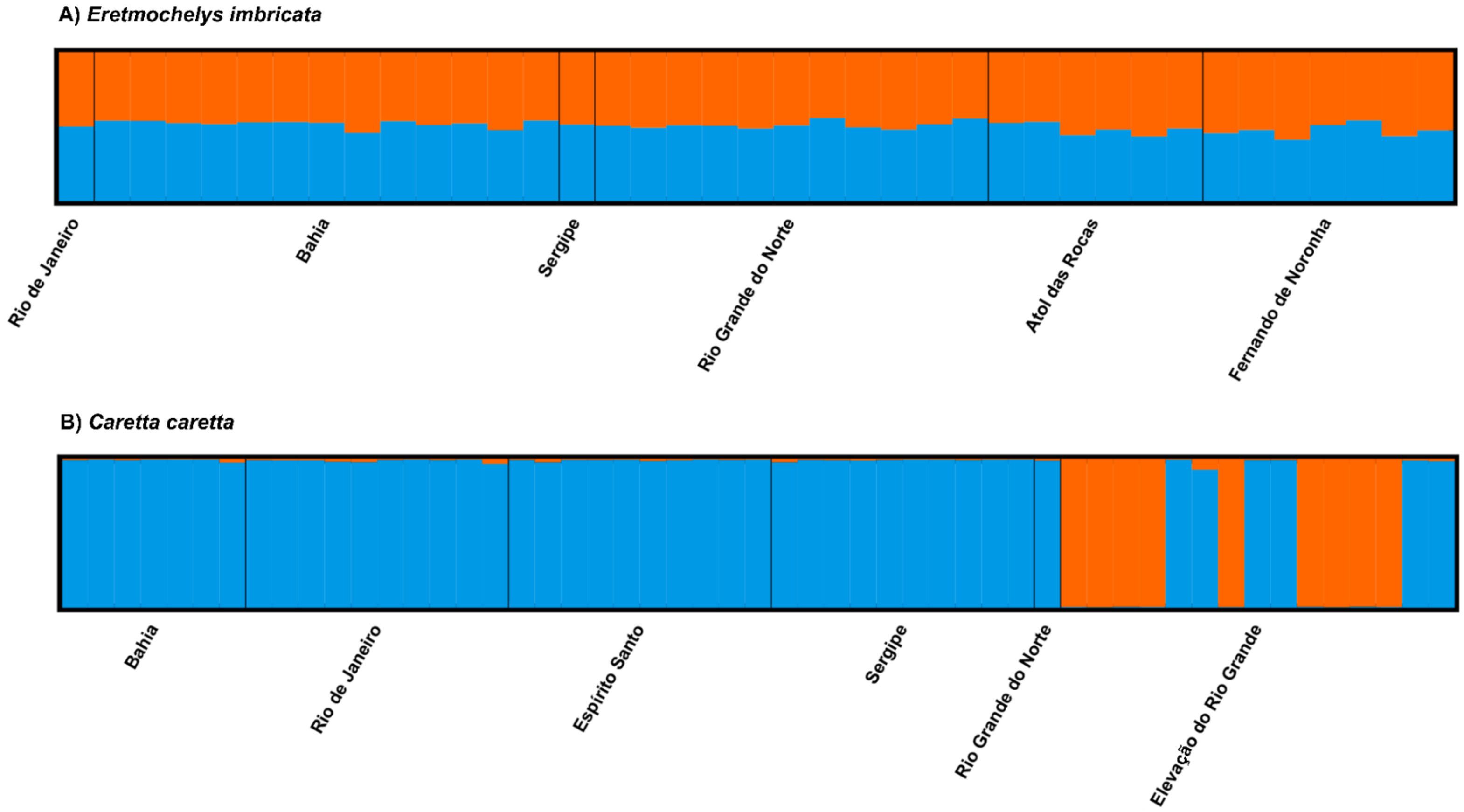


Figure S1. STRUCTURE bar plots representing K = 2 using correlated allele frequencies model for *Eretmochelys imbricata* and *Caretta caretta* from different Brazilian populations. The x-axis represents each individual analyzed and the y-axis represents the estimated admixture proportions related to each population. This graphic was obtained with CLUMPAK.
