## Supplementary Figure S2 for "New genetic insights about hybridization and population structure of hawksbill and loggerhead turtles from Brazil"

**Arantes_SupMat_FigureS2_Hybridization of Brazilian sea turtles**

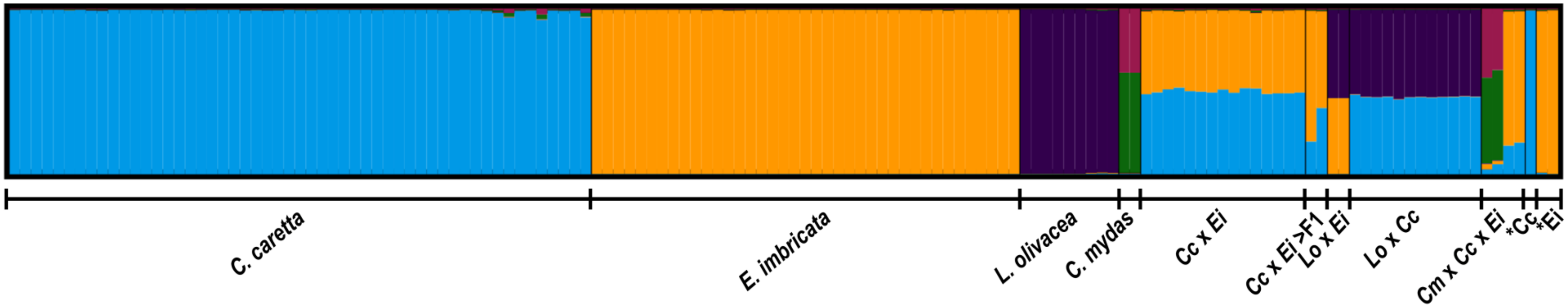


Figure S2. STRUCTURE bar plots representing K = 5 using independent allele frequencies model. The x-axis represents each individual analyzed and the y-axis represents the estimated admixture proportions related to each parental species. This graphic was obtained with CLUMPAK. The asterisks (*) show misidentified individuals. Cc: *Caretta caretta*, Ei: *Eretmochelys imbricata*, Lo: *Lepidochelys olivacea*, Cm: *Chelonia mydas*, >F1: introgressed hybrid.
