## Supplementary Figure S3 for "New genetic insights about hybridization and population structure of hawksbill and loggerhead turtles from Brazil"

**Arantes_SupMat_FigureS3_Hybridization of Brazilian sea turtles**

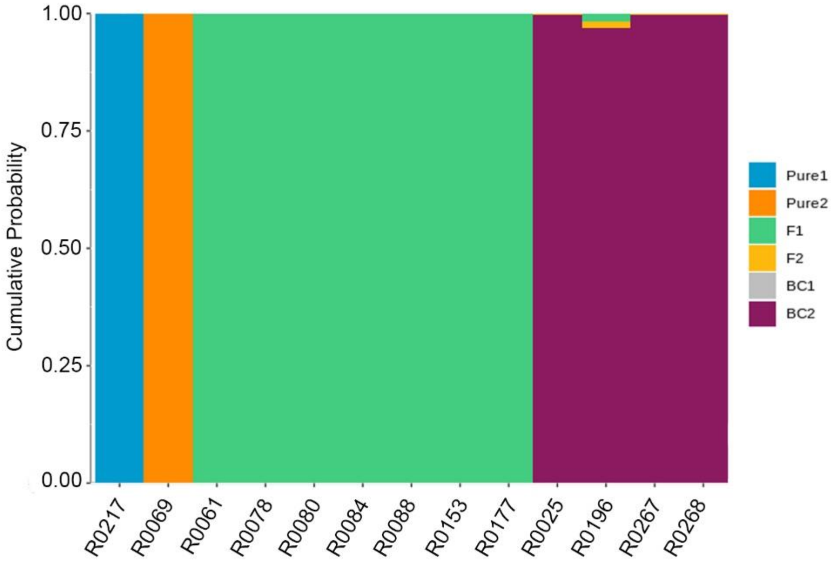


Figure S3. NewHybrids analysis of 13 individuals previously identified as introgressed hybrids. Each vertical bar represents one individual and the y-axis represents its posterior probability of belonging to different classes: *C. caretta* (Pure 1), *E. imbricata* (Pure 2), F1 hybrid, F2 hybrid, backcross with *C. caretta* (BC1) or backcross with *E. imbricata* (BC2). This graphic was obtained with R package HybridDetective.
